# Depressive-like behaviors induced by somatostatin-positive GABA neuron silencing are rescued by alpha 5 GABA-A receptor potentiation

**DOI:** 10.1101/2020.10.05.326306

**Authors:** Corey Fee, Thomas D. Prevot, Keith Misquitta, Daniel E. Knutson, Guanguan Li, Prithu Mondal, James M. Cook, Mounira Banasr, Etienne Sibille

**Author notes:** Corresponding author, Etienne Sibille, PhD, CAMH, 250 College Street, Room 134, Toronto, ON M5T 1R8, Canada.

## Abstract

**Introduction:** Deficits in somatostatin-positive gamma-aminobutyric acid interneurons (“SST+ cells”) are associated with major depressive disorder (MDD) and a causal link between SST+ cell dysfunction and depressive-like deficits has been proposed, based on rodent studies showing that chronic stress induces a low SST+ GABA cellular phenotype across corticolimbic brain regions, that lowering *Sst*, SST+ cell, or GABA functions induces depressive-like behaviors, and that disinhibiting SST+ cell functions has antidepressant effects. Recent studies found that compounds preferentially potentiating receptors mediating SST+ cell functions with α5-GABA-A receptor positive allosteric modulators (α5-PAMs) achieved antidepressant-like effects. Together, evidence suggests that SST+ cells regulate mood and cognitive functions that are disrupted in MDD and that rescuing SST+ cell function may represent a promising therapeutic strategy.

**Methods:** We developed a mouse model with chemogenetic silencing of brain-wide SST+ cells and employed behavioral characterization 30 min after acute or sub-chronic silencing to identify contributions to behaviors related to MDD. We then assessed whether an α5-PAM, GL-II-73, could rescue behavioral deficits induced by SST+ cell silencing.

**Results:** Brain-wide SST+ cell silencing induced features of stress-related illnesses, including elevated neuronal activity and plasma corticosterone levels, increased anxiety- and anhedonia-like behaviors, and impaired short-term memory. GL-II-73 led to antidepressant-like improvements among all behavioral deficits induced by brain-wide SST+ cell silencing.

**Conclusion:** Our data validate SST+ cells as regulators of mood and cognitive functions, support a role for SST+ cell deficits in depressive-like behaviors, and demonstrate that bypassing low SST+ cell function via α5-PAM represents a targeted antidepressant strategy.

**Significance Statement:** Human and animal studies demonstrate somatostatin-positive GABAergic interneuron (“SST+ cell”) deficits as contributing factors to the pathology of major depressive disorder (MDD). These changes involve reduced SST and GABAergic markers, occurring across corticolimbic brain regions. Studies have identified roles for SST+ cells in regulating mood and cognitive functions, but employed genetic or region-specific ablation that is not representative of disease-related processes. Here, we developed a chemogenetic mouse model of brain-wide low SST+ cell function. This model confirmed a role for SST+ cells in regulating anxiety- and anhedonia-like behaviors, overall behavioral emotionality, and impaired working memory. We next showed that a positive allosteric modulator at α5-GABA-A receptors (α5-PAM, GL-II-73) rescued behavioral deficits induced by low SST+ cell function. These findings support a central role for brain-wide low SST+ cell function in MDD and validate targeting α5-GABA-A receptors as a therapeutic modality across MDD symptom dimensions.

## Introduction

Major depressive disorder (MDD) is a severe psychiatric illness affecting 322 million people, a leading cause of disability worldwide (Friedrich, 2017; Vos et al., 2017; World Health Organization, 2017; Rehm and Shield, 2019) and is more common among females (5.1%) than males (3.6%) (World Health Organization, 2017). The diagnosis and treatment of MDD is limited by heterogeneity in pathological and clinical presentation, with low mood, anhedonia, physiological and cognitive impairments, and highly comorbid anxiety symptoms. Further, current first-line monoaminergic antidepressants are ineffective in 50% of treated patients (Gaynes et al., 2009; Holtzheimer and Mayberg, 2011), highlighting a need to characterize contributing pathologies to develop targeted drug therapies that reverse or circumvent cellular deficits. Four decades of human and animal studies demonstrate reduced level, synthesis, and function of neurons expressing the inhibitory neurotransmitter, gamma-aminobutyric acid (GABA), associated with MDD (Luscher et al., 2011; Newton et al., 2019). Among these, GABA neurons co-expressing the neuropeptide somatostatin (“SST+ GABA cells” or “SST+ cells”) are selectively vulnerable in MDD (Lin and Sibille, 2013; Fee et al., 2017).

GABA neurons play a critical role in information processing by cortical microcircuits to regulate mood and cognitive functions. Cortical microcircuits encode neuronal information through activity patterns of excitatory glutamatergic pyramidal neuron (PN) ensembles. GABA neurons shape this activity differently based on non-overlapping populations that co-express SST, the calcium-binding protein parvalbumin (PV+), or the ionotropic serotonin receptor 5HT3aR with or without vasoactive intestinal peptide (VIP+) (Rudy et al., 2011). SST+ cells gate excitatory PN input via dendritic inhibition and modulate output via perisomatic disinhibition through SST-PV+ cell afferents (Fee et al., 2017; Hu et al., 2019). Dendritic SST+ cell inhibition is partially mediated by GABA acting on GABA-A and GABA-B receptors (GABAA-R/GABAB-R) (Urban-Ciecko et al., 2015; Schulz et al., 2018), whereas the SST peptide is released under distinct firing conditions (Katona et al., 2014; Yavorska and Wehr, 2016). SST has shared roles alongside GABA in pre- and post-synaptic PN inhibition (Tallent and Siggins, 1997; Schweitzer et al., 1998), plus distinct roles in negative regulation of stress-related hypothalamic-pituitary adrenal (HPA) axis activation (Stengel and Taché, 2017).

Histological studies in postmortem MDD subjects identified reductions of SST and GABA markers at the mRNA and protein level in the prefrontal cortex (PFC) (Sibille et al., 2011), anterior cingulate cortex (ACC) (Tripp et al., 2011), and amygdala (Guilloux et al., 2012) that were more severe in females (Seney et al., 2013; Seney, M., Tripp, A., McCune, S., Lewis, D., Sibille, 2015). Follow-up studies indicated reduced SST mRNA per cell, spanning all ACC layers (Tripp et al., 2011), and reduced detectable SST+ cell density without overall cell density changes in the amygdala (Douillard-Guilloux et al., 2016). *In vivo* studies also revealed MDD-related reductions in PFC (Hasler et al., 2007) and ACC (Price et al., 2009) GABA levels, with reduced GABAA-R and GABAB-R neurotransmission (Bajbouj et al., 2006; Levinson et al., 2010; Radhu et al., 2013). The regional and functional differentiation of GABAA-Rs depend upon heteropentameric subunit composition, commonly containing two α, two β, and one γ-subunit (Rudolph et al., 2001; Sieghart and Sperk, 2002). SST+ cell functions are partially mediated by α5 subunit-containing-GABAA-Rs given their localization to PN dendrites and PV+ cells in the PFC, hippocampus (HPC), and ventral striatum (Ali and Thomson, 2008; Schulz et al., 2018; Hu et al., 2019). *GABRA5* transcripts, encoding α5GABAA-Rs, were also reduced in the PFC of MDD and aged subjects (Oh et al., 2019). Together, this evidence suggests that MDD involves a low SST+ GABA cellular phenotype affecting pre- and post-synaptic functions and arising from intrinsic cellular vulnerability, rather than layer- or region-specific origins.

Preclinical studies support a role for SST + cells in regulating mood and cognitive functions, and suggest that disrupted SST+ cell function may contribute to deficits reflecting MDD symptom dimensions. Rodent chronic stress studies found reduced mRNA of *Sst* and the GABA-synthesizing enzyme *Gad67* in the cingulate cortex, and selectively altered SST+ cell and not PN transcripts, linked to disrupted growth factor signaling and cellular stress (Lin and Sibille, 2015; Girgenti et al., 2019). Studies in mice with genetic *Sst* deletion recapitulated elevated corticosterone, reduced growth factor and *Gad67* expression, and elevated depressive-like behaviors (Lin and Sibille, 2015). Furthermore, acute chemogenetic or optogenetic PFC SST+ cell silencing induced depressive-like behaviors (Soumier and Sibille, 2014) and working memory impairment (Abbas et al., 2018), whereas genetic disinhibition had antidepressant-like effects (Fuchs et al., 2016). These results implicate SST+ cells as key regulators of mood and cognitive functions; however, genetic ablation or region-specific disruptions may not recapitulate disease-related changes observed in humans (Fee et al., 2017).

Functional neuroimaging studies demonstrated normalization of GABA levels after antidepressant treatment across pharmacological, cognitive-behavioral, and neuromodulatory modalities (Fee et al., 2017). Although first-line antidepressants do not target GABA or SST deficits directly, mixed antidepressant efficacy has been observed from monotherapy or combination therapy with benzodiazepines (BZDs) (Benasi et al., 2018; Gomez et al., 2018; Ogawa et al., 2019) that act as non-selective positive allosteric modulators (PAMs) at GABAA-Rs between γ2 and α1-3, 5 subunits (Gomez et al., 2018). However, BZDs have side effects and abuse liability attributed to their pan-α-subunit selectivity (Vgontzas et al., 1995). Recent preclinical studies assessing BZD-derivatives with selectivity for α5-GABAA-Rs (α5-PAMs), which have restricted corticolimbic distribution, found anxiolytic-, antidepressant-like, and pro-cognitive effects in young, aged, and chronic stress-exposed rodents (Koh et al., 2013; Piantadosi et al., 2016; Prevot et al., 2019b). α5-PAMs may optimize α-subunit selectivity, wherein for instance, α1 was associated with sedative, amnesic, and addictive properties, α2/3 with anxiolysis, and α5 with mood and cognitive regulation (Prevot and Sibille, 2020).

Based on the cumulative evidence for low SST+ cell function as a contributing pathology in MDD, we aimed to determine whether brain-wide low SST+ cell function induced behavioral deficits related to MDD symptom dimensions: anxiety- and anhedonia-like behavior, behavioral emotionality, and impaired memory. We employed behavioral characterization of mice with repeated acute chemogenetic brain-wide SST+ cell function silenced. We next determined whether an α5-selective PAM, GL-II-73 (Prevot et al., 2019b), could rescue behavioral deficits. We predicted that brain-wide SST+ cell silencing induces depressive-like behavioral deficits that can be rescued by α5-PAM.

## Materials and Methods

A complete description of methods/study designs can be found in <u>Supplementary Information</u>.

### Animals

Viral transduction was validated in C57BL/6J *Sst^Gfp/+^* mice (Taniguchi et al., 2011; Lin and Sibille, 2015). Behavioral and molecular experiments were performed using C57BL/6J *Sst^Cre/+^* mice (*Sst^tm2.1(cre)Zjh^*/J, Jackson Laboratories, ME; #913944), postnatal day zero (PND0) at surgery and 9-16W at testing. Mice were maintained under 12-hour light/dark cycle (7am-7pm) with food and water *ad libitum*, group-housed (4/cage), except during behavioral testing (1/cage). Tests were performed according to Institutional and Canadian Council on Animal Care (CCAC) guidelines.

### AAV Vectors and Neonatal Injection

Enhanced serotype inhibitory Designer Receptors Exclusively Activated by Designer Drugs (DREADD) vectors used were AAV-PHP.eB-hSyn-DIO-hM4D(Gi)-mCherry (Addgene #44362, ME) (Chan et al., 2017). Control vectors used were AAV-PHP.eB-hSyn-DIO-mCherry (Addgene #50459).

Brain-wide SST+ cell-specific hM4Di expression was achieved by low-dose bilateral intracerebroventricular (i.c.v.) infusion of PHP.eB serotype Flip-Excision (FLEx-ed) AAVs in PND0 *Sst^Cre/+^* mice (Kim et al., 2013) (**Supplementary Methods**). For validation experiments, *Sst^Gfp/+^* neonates received DREADD vectors. For characterization experiments, *Sst^Cre/+^* neonates received DREADD (*Sst^hSyn-hM4Di-mCherry^* mice) or Control (*Sst^hSyn-mCherry^* mice) vectors.

### Chemogenetic Inhibition of Brain-wide SST+ cells & α5-PAM Co-administration

For the first characterization experiment, *Sst^hSyn-hM4Di-mCherry^* mice received clozapine-N-oxide (CNO=3.5mg/kg) or vehicle (Veh=0.9% saline) intraperitoneally (i.p.) 30 min before testing (*n*=16/group; 50% female; **Fig 2*a***). We confirmed that locomotor activity and anxiety-like behavior were not affected by CNO in separate *Sst^Cre/+^* cohorts not expressing hM4Di (**Fig S1**).

For the second characterization experiment, we assessed whether deficits induced by SST+ cell silencing could be rescued by co-administering CNO and GL-II-73 (Prevot et al., 2019b). *Sst^hSyn-mCherry^* and *Sst^hSyn-hM4Di-mCherry^* mice were generated and all groups received 3.5mg/kg CNO, achieving SST+ cell silencing only in hM4Di-expressing mice (“SST-silenced”=*Sst^hSyn-hM4Di-mCherry^*, “SST-control”=*Sst^hSyn-mCherry^*). Mice were randomly assigned to receive Veh (vehicle+CNO) or 10mg/kg GL-II-73 (+vehicle+CNO) totaling 4 groups (*n*=10-12/group; 50% female; **Fig 4*a***).

### Behavioral Analyses

In adulthood, *anxiety-like behavior* was assessed with the PhenoTyper test (PT), elevated plusmaze (EPM), open field test (OFT) and novelty-suppressed feeding test (NSF), *anhedonia-like behavior* with sucrose consumption test (SCT), *mixed anxiety-/anhedonia-like behavior* with novelty-induced hypophagia test (NIH), *antidepressant-predictive behavior* in the forced-swim test (FST), and *memory impairment* in the Y-maze (YM) and novel object recognition test (NORT), following past study designs (Fee et al., 2020). CNO was administered 30 min prior to test initiation, except for the PT where CNO was administered 30 min prior to the dark cycle (90 min prior to light challenge) to allow habituation.

Z-score normalization captured changes along behavioral dimensions, directionally, relative to controls (i.e., increase = deficit, decrease = improvement) for anxiety- (PT, EPM, OFT, NSF, NIH) and anhedonia-like behavior (NIH, SCT), overall behavioral emotionality (all previous + FST), and working- and short-term memory impairment (YM, NORT) (Guilloux et al., 2011) (**Supplementary Methods**). Viral transduction efficiency and SST+ cell specificity was visually inspected via immunohistochemistry (IHC) in all mice following behavioral experiments.

### Immunohistochemistry and Microscopy

The transduction efficiency and SST+ cell specificity of neonatally-delivered i.c.v. AAV-PHP.eB-hSyn-DIO-hM4D(Gi)-mCherry was assessed via IHC in sections from paraformaldehyde-perfused *Sst^Gfp/+^* mice fluorescently labeled for GFP and mCherry (*n=*8, 2 fields/region/mouse; **Fig 1*a***).

**Figure 1.**
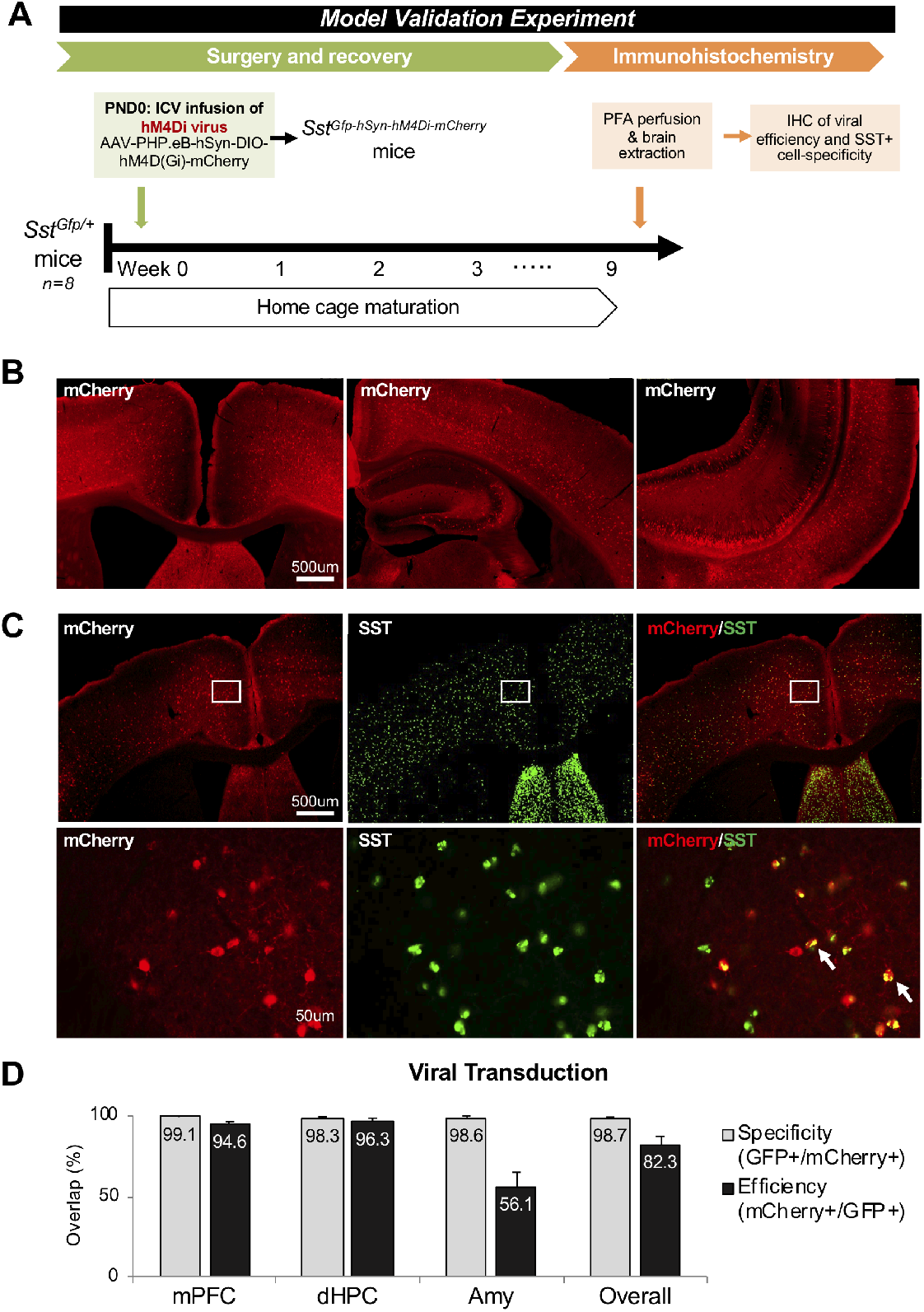
Validation of brain-wide SST+ cell targeting. (**A**) Experimental design for validation of SST+ cell targeting by 2×10^10^vg AAV-PHP.eB-DIO-hM4Di-mCherry i.c.v. in PND0 *Sst^Gfp/+^* mice, validated by immunohistochemistry (IHC). (**B**) Representative images of brain-wide viral transduction in adult mice (red = conditional mCherry expression; scale = 500 μm). (**C**) Representative images of hM4Di-mCherry (red) and SST-GFP (green) co-expression (white arrow) in the mPFC. (**D**) Efficiency (mCherry/GFP) and specificity (GFP/mCherry) of viral transduction in mPFC, dHPC, and Amy (*n*=8 mice).

Forty-eight hours after the last test, *Sst^hSyn-hM4Di-mCherry^* mice from behavioral experiments were injected with Veh or CNO and perfused 110 minutes later (approximating CNO + c-Fos peaks) to quantify neuronal activation following SST+ cell silencing, indexed by c-Fos+ cell counts, (Dragunow and Faull, 1989; Ferguson et al., 2011; Lin and Sibille, 2015) (**Fig 2*a***). Cell counts were collected in every third section (N=4-5 sections/region/mouse) for medial PFC (mPFC), dorsal hippocampus (dHPC), and basolateral amygdala (Amy). Image processing and counting was performed under blinded conditions for Veh/CNO groups (*n*=8/group) using Fiji (Schindelin et al., 2012).

**Figure 2.**
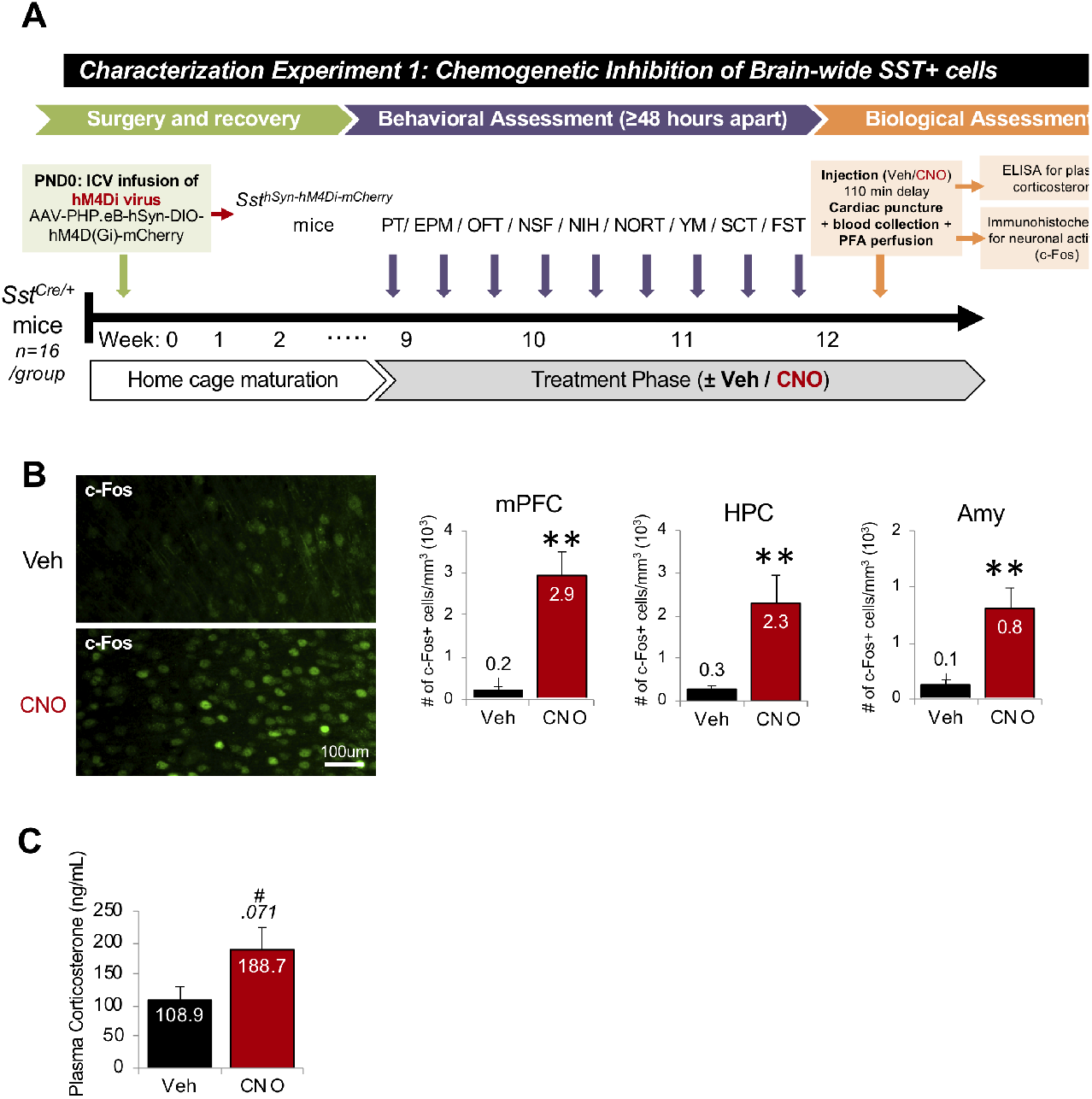
Validation of brain-wide SST+ cell manipulation. (**A**) Experimental design for behavioral and biological characterization following chemogenetic SST+ cell silencing in *Sst^hSyn-hM4Di-mCherry^* mice. (**B**) Representative image of c-Fos+ cells 110 min after treatment with vehicle (Veh, top) or 3.5 mg/kg clozapine-N-oxide (CNO, bottom). CNO-hM4Di activation increased neuronal activity (c-Fos+ cell counts) in the mPFC, HPC, and amygdala (Amy) (*n*=8 mice/group). (**C**) From the same group, CNO induced a trend elevation in plasma corticosterone (*n*=16 mice/group). \*\**p*<.01; ^#^*p*<.1.

### Corticosterone Measurement

In *Sst^hSyn-hM4Di-mCherry^* mice from behavior & c-Fos experiments, plasma corticosterone was assessed by ELISA (Arbor Assays, MI) 110 minutes after Veh/CNO administration from blood collected via cardiac puncture prior to perfusions (*n*=16 mice/group).

### Data Analysis

Data were analyzed using SPSS (IBM, NY) and expressed as mean ± standard error of the mean (SEM). For Veh vs. CNO, data were analyzed using one-way analysis of covariance (ANCOVA) with treatment as independent variable and sex as covariate. For SST-control vs. SST-silenced, data were analyzed using two-way ANCOVA with group and treatment (Veh vs. GL-II-73) as independent variables and sex as covariate, using Bonferroni-adjusted *post hoc* where appropriate. Timecourse parameters were assessed by repeated-measures ANCOVA with Greenhouse-Geisser correction. Object recall in the NORT was assessed using paired-samples *t*-tests.

## Results

### Intracerebroventricular AAV-PHP.eB DREADD infusion in neonatal *Sst^Cre^* mice enables brain-wide manipulation of SST+ cells in adulthood

Validation of SST+ cell viral targeting was performed in *Sst^Gfp/+^* mice administered low-dose (2×10^10^vg) AAV-PHP.eB-DIO-hM4Di-mCherry via neonatal i.c.v. infusion (Soumier and Sibille, 2014; Lin and Sibille, 2015) (**Fig 1*a-d***). In adults, IHC quantification revealed high viral transduction efficiency (mPFC=95±2%, dHPC=96±2%, Amy=56±9%, Overall=82±5%) and SST+ cell specificity (mPFC=99±1%, dHPC=98±1%, Amy=99±1%, Overall=99±1%) that was consistent with *Sst* expression patterns (NCBI, 2020) (**Fig 1*b-d***).

Based on established roles for GABA and SST in inhibitory regulation of local neuronal activity and endocrine signaling (Yavorska and Wehr, 2016), chemogenetic SST+ cell silencing was next validated in a separate cohort of *Sst^hSyn-hM4Di-mCherry^* mice by quantifying neuronal activity (via c-Fos+ cell counts (Kovács, 1998)) and plasma corticosterone levels 110 minutes following SST+ cell inactivation by CNO (**Fig 2*a***). Compared to Veh, CNO increased the number of c-Fos+ cells in the mPFC (*F_1,13_*=19.42; *p*=.001), HPC (*F_1,13_*=14.05; *p*=.002), and Amy (*F_1,13_*=19.35; *p*=.001; **Fig 2*b***). CNO induced a trend-level elevation in corticosterone as assessed by ELISA (*F_1,28_*=3.51; *p*=.071; **Fig 2*c***). Sex did not influence c-Fos or corticosterone levels.

### Brain-wide SST+ cell silencing increases anxiety-like behaviors, overall behavioral emotionality, and memory impairment

In the characterization experiment 1 cohort, we sought to determine whether SST+ cell silencing induced anxiety-like behavior (PT, EPM, OFT, NSF), anhedonia-like behavior (NIH, SCT), antidepressant-predictive behavior (FST), and memory impairment (YM, NORT).

Analysis of PT shelter zone activity before, during, and after a 1h light challenge in the dark cycle revealed significant main effects of time (*F_7.51,217.82_*=2.42;*p*< .05), group (*F_1,29_*=6.75; *p*<.05), and group*time interaction (*F_7.51,217.82_*=2.18; *p*<.05; **Fig 3*a***). CNO significantly increased shelter zone time after injection and persisting for 6 hours after light challenge (*p*<.05). Overall anxiogenic response, indexed by shelter zone time area under the curve (AUC) from light challenge initiation until test completion confirmed that CNO significantly increased shelter zone time (*F_1,29_*=11.97; *p*<.01; **Fig 3*b***).

**Figure 3.**
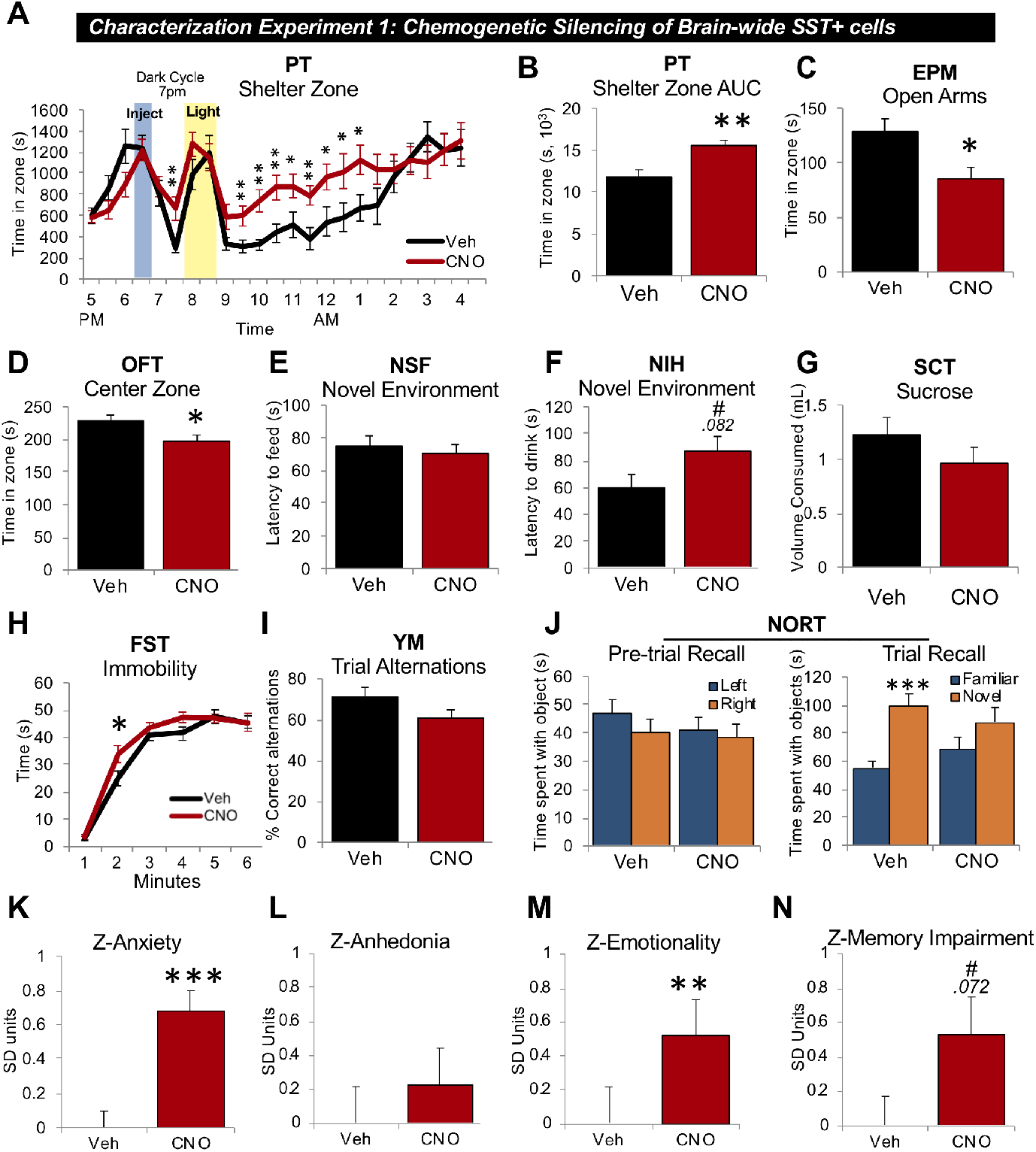
Brain-wide SST+ cell silencing in *Sst^hSyn-hM4Di-mCherry^* mice elevates anxiety-like behavior, overall behavioral emotionality, and memory impairment. (**A-J**) Behavioral tests administered 30 minutes after injection of vehicle (Veh) or 3.5mg/kg clozapine-N-oxide (CNO) i.p. except for the PhenoTyper Test (PT) (administered 90 min prior to light challenge). (**A**) Shelter zone time before, during and after light challenge in the PT. (**B**) Summary area under the curve (AUC) of shelter zone time from light challenge initiation until the test completion. (**C**) Time spent in open arms of the elevated plus maze (EPM). (**D**) Time spent in center zone of the open field test (OFT). (**E**) Novel environment latency to feed in the novelty suppressed feeding (NSF) test. (**F**) Novel environment latency to drink milk reward in the novelty-induced hypophagia test (NIH). (**G**) Sucrose consumed in the sucrose consumption test (SCT). (**H**) Immobility time in the forced swim test (FST). (**I**) Percent correct alternations in Y-maze (YM). (**J**) Pre-trial left/right familiar object recall (left) and trial familiar/novel object recall (right). (**K-N**) Z-scores integrating test parameters assessing anxiety-like behavior (**K**), anhedonia-like behavior (**L**), behavioral emotionality (**M**), and memory impairment (**N**). *n*=16 mice/group; 50% male/female. \*\*\**p*<.001; \*\**p*<.01; \**p*<.05; ^#^*p*<.01. Sex effects and homecage measures in Table S1.

Anxiogenic CNO effects were also detected from decreased EPM open arm time (*F_1,25_*=7.06; *p*<.05; **Fig 3*c***) and OFT center zone time (*F_1,29_*=5.86; *p*<.05; **Fig 3*d***). Distance travelled was unchanged between groups in the OFT and EPM (**Table S1**). In the NSF, no group differences were detected for novel environment latency to feed (**Fig 3*e***). In the NIH, CNO induced a trend-level increase in novel environment latency to drink (*F_1,27_*=3.26; *p*=.082; **Fig 3*f***). Locomotor activity (PT, EPM, OFT) and home cage latencies to feed or drink (NSF/NIH) did not differ between groups (**Table S1**). There were no effects of sex for PT, EPM, OFT, NSF, or NIH parameters.

In the SCT, no group differences were detected for sucrose consumed (**Fig 3*g***), or sucrose ratio [sucrose/(sucrose + water consumed)]; **Table S1**). Covariate analyses revealed that females consumed significantly less sucrose (*F_1,28_*=7.7; *p*<.01; **Table S1**).

FST analysis revealed a significant effect of time (*F_3.46, 100..39_*=15.15; *p*< .001) and trendlevel time*treatment interaction (*F_3.46,100.39_*=2.32; *p*=.071) as CNO increased immobility in minute two (*p*<.05; **Fig 3*h***).

In the YM, group differences were not detected for pre-trial (**Table S1**) or trial alternation rates (**Fig 3*i***). In the NORT pre-trial, left/right familiar object times were equivalent for both groups (**Fig 3*j***). In the NORT trial, Veh-treated mice spent significantly more time with the novel vs. familiar object (*t*=4.44, df=15; *p*<.001), while CNO-treated mice did not (*t*=1.56; df=15; *p*=.14), indicating impaired short-term recall. There was no effect of sex for FST, YM, or NORT parameters.

Given the well-characterized variability associated with preclinical behavioral tests (Willner, 2017; Prevot et al., 2019c), we next used Z-scores to assess the consistency of behavioral responses across tests assessing *a priori* related dimensions (Guilloux et al., 2011). CNO significantly increased Z-anxiety scores, reflecting PT, EPM, OFT, NSF and NIH parameters (*F_1,29_*=17.53; *p*<.001; **Fig 3*k***). Z-anhedonia scores did not differ between groups, reflecting SCT and NIH (**Fig 3*l***). CNO significantly elevated Z-emotionality scores, reflecting all previous tests plus FST (*F_1,29_*=13.96; *p*<.01; **Fig 3*m***). CNO induced a trend-level increase in Z-memory impairment scores, reflecting YM and NORT (*F_1,29_*=3.49; *p*=.072; **Fig 3*n***).

### α5-PAM (GL-II-73) rescues behavioral deficits induced by brain-wide SST+ cell silencing

SST-control (*Sst^hSyn-mCherry^*) and SST-silenced (*Sst^hSyn-hM4Di-mCherry^*) mice were generated by neonatal infusion of Control or DREADD viruses in *Sst^Cre/+^* mice, and in adulthood administered Veh (vehicle+CNO) or GL-II-73 (vehicle+CNO+GL-II-73) prior to behavioral testing (**Fig 4*a***).

**Figure 4.**
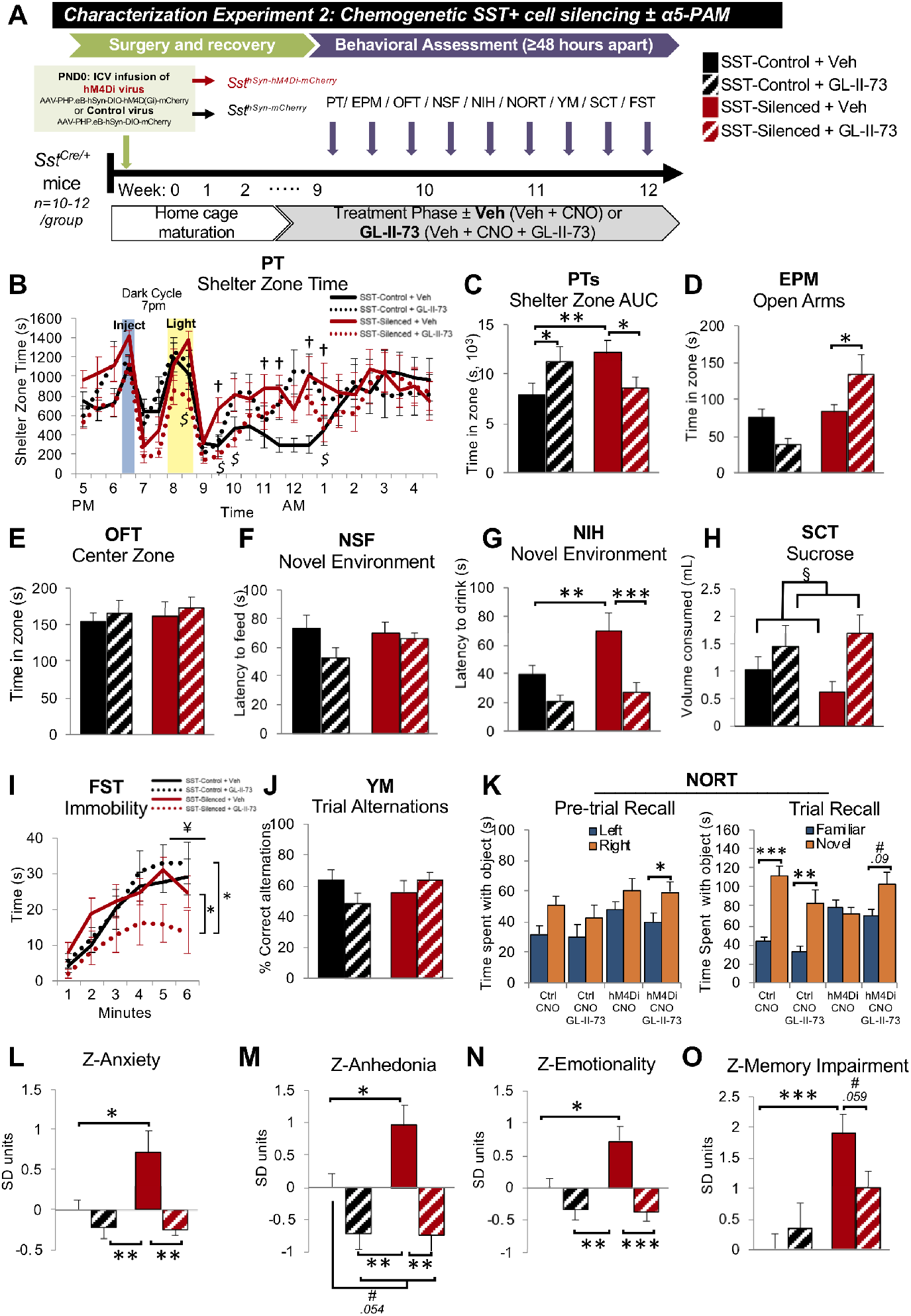
GL-II-73 rescues behavioral deficits induced by brain-wide SST+ cell silencing. (A-J) Behavioral tests administered 30 minutes following i.p. injection of vehicle + CNO (Veh) or vehicle + CNO + GL-II-73 (GL-II-73) in SST-control (*Sst^hSyn-mCherry^*) or SST-silenced (*Sst^hSyn-hM4Di-mCherry^*) mice (except in PhenoTyper as indicated). (**B**) Shelter zone time before, during, and after light challenge and (**C**) derived shelter zone area under the curve (AUC) from light challenge initiation until PhenoTyper test (PT) completion. (**D**) Time spent in open arms of the elevated plus maze (EPM). (**E**) Time spent in center zone of the open field test (OFT). (**F**) Novel environment latency to feed for the novelty suppressed feeding (NSF) test. (**G**) Novel environment latency to drink milk reward for the novelty-induced hypophagia test (NIH). (**H**) Sucrose consumed in the sucrose consumption test (SCT). (**I**) Immobility time in the forced swim test (FST). (**J**) Percent correct alternations in Y-maze (YM). (**K**) Pre-trial left/right familiar object recall (left) and trial familiar/novel object recall (right) in the novel object recognition test (NORT). (**L-O**) Z-scores integrating test parameters assessing anxiety-like behavior (**L**), anhedonia-like behavior (**M**), overall behavioral emotionality (**N**), and memory impairment (**O**). *n* = 16 mice/group; 50% male/female. \*\*\**p*<.001; \*\**p*<.01; \**p*<.05; ^#^*p*<.01; †=*p*<.05 SST-Control+Veh vs. SST-Silenced+Veh; $=*p*<.05 SST-Silenced+GL-II-73 vs. SST-Silenced+Veh; § = significant main effect of treatment; ¥ = significant main effect of group. Sex effects and homecage parameters in Table S2.

Analysis of PT shelter zone time revealed a significant main effect of time (*F_9.5,389.4_*=6.971; *p*<.001) and time*group*treatment interaction (*F_9.5,389.4_*=2.1; *p*<.05), wherein SST-silenced+Veh mice had significantly increased shelter zone time after the light challenge (*p*<.05 vs. SST-control+Veh; **Fig 1*b***). This effect was reversed by GL-II-73 in SST-silenced mice (*p*<.05). For shelter zone AUC, ANOVA revealed a group*treatment interaction (*F_1,41_*=8.45; *p*<.01), wherein SST-silenced+Veh mice had increased AUC (*p*<.01 vs. SST-control+Veh) that was similarly reversed by GL-II-73 treatment (*p*<.05; **Fig 4*c***). Increased shelter zone AUC was also detected in SST-control+GL-II-73 vs. SST-control+Veh mice (*p*<.05). Covariate analyses revealed that males had significantly increased shelter zone AUCs (*F_1,41_*=22.83; *p*<.001; **Table S2**). Locomotor activity did not differ between groups (*F_1,41_*=.65; *p*>.59; **Table S2**).

Analysis of EPM open arm time revealed a significant main effect of group (*F_1,35_*=11.63; *p*=.002) and group*treatment interaction (*F_1,35_*=8.83; *p*=.005) (**Fig 4*d***). GL-II-73 increased open arm time in SST-silenced mice (*p*<.05 vs. SST-silenced+Veh). OFT center zone time was unchanged between groups (**Fig 4*e***). However, CNO+GL-II-73 significantly increased distance travelled in EPM (*p*=.044) and OFT (*p*<.001) (**Table S2**). In the NSF, no group differences were detected for novel environment latency to feed (**Fig 4*f***). Analyses of NIH novel environment latency to drink revealed significant main effects of group (*F_1,40_*=5.31; *p*<.05), treatment (*F_1,40_*=16.66; *p*<.001), and trend-level group*treatment interaction (*F_1,40_*=2.93; *p*=.095), wherein SST-silenced+Veh mice had increased latency (*p*<.01 vs. SST-control+Veh) that was rescued by GL-II-73 (*p*<.05) (**Fig 4*g***). Covariate analysis revealed that females had significantly increased latency to drink (*F_1,40_*=6.57; *p*<.05; **Table S2**). Home cage latencies to feed (NSF) or drink (NIH) did not differ between groups (**Table S2**).

In the SCT, a significant main effect of treatment was detected; GL-II-73 increased sucrose consumption (*F_1,38_*=6.25; *p*<.05; **Fig 4*h***), but not sucrose ratio (**Table S2**), indicating increased overall fluid consumption rather than preference.

In the FST, a significant main effect of time (*F_3.19,102_*=6.74; *p*<.001), time*group interaction (*F_3.19,102_*=3.2; *p*<.05), and group*treatment interaction were detected (*F_1,32_*=6.87; *p*<.05; **Fig 4*i***). GL-II-73 decreased immobility in SST-silenced mice (*p*<.05 vs. SST-silenced+Veh, SST-control+GL-II-73). However, this may have been confounded by the previously identified hyperlocomotor effect of GL-II-73.

In the YM, pre-trial (**Table S2**) and trial (**Fig 4*j***) alternation rates did not differ between groups. In the NORT pre-trial, only SST-silenced+GL-II-73 mice spent significantly more time with the right-located object, indicating side preference that was balanced by rotating novel object position in the trial (*t*=2.84, df=10; *p*=.018; **Fig 4*k***). In the NORT trial, significant short-term recall was detected for SST-control+Veh (*t*=1.62, df=11; *p*<.001) and SST-control+GL-II-73 (*t*=3.89, df=10; *p*=.003). SST+ cell silencing impaired familiar object recall (*t*=.695, df=10; *p*=.503) that was partially rescued by GL-II-73 (*t*=1.85, df=10; *p*=.09).

For Z-score analysis, EPM, OFT (Z-anxiety), and FST parameters (Z-emotionality) were excluded due to potential locomotor bias, whereas this effect was not detected in other tests. Analysis of Z-anxiety scores (PT, NSF, NIH) revealed a significant main effect of treatment (*F_1,41_*=11.18; *p*<.01), a trend-level group effect (*F_1,41_*=3.94; *p*=.054), and a significant group*treatment interaction (*F_1,41_*=4.32; *p*<.05) (**Fig 4*l***). SST-silenced+Veh mice had elevated anxiety-like behavior (*p*<.05 vs. SST-control+Veh). GL-II-73 had anxiolytic effects in SST-control+GL-II-73 mice and reversed elevated anxiety-like behavior in SST-silenced+GL-II-73 mice (*ps*<.01 relative to SST-silenced+Veh; **Fig 4*l***). Z-anhedonia scores (NIH, SCT), ANCOVA detected significant treatment effects (*F_1,41_*=20.13; *p*<.001), and trend-level group (*F_1, 41_*=3.34; *p*<.075) and group*treatment effects (*F_1,41_*=3.8; *p*=.056). Z-anhedonia was significantly elevated in SST-silenced+Veh mice (*p*<.05 vs. SST-control+Veh) and reduced or rescued in SST-Control+GL-II-73 and SST-Silenced+GL-II-73 mice, respectivey (*ps*<.01 relative to SST-silenced+Veh; **Fig 4*m***). Z-emotionality scores (all previous tests) were significantly affected by treatment (*F_1,41_*=17.5; *p*<.001), trend-level group effects (*F_1,41_*=4.1; *p*=.05), and a significant group*treatment interaction (*F_1,41_*=4.77; *p*<.05). Z-emotionality scores were increased in SST-silenced+Veh mice (*p*<.05 vs. SST-control+Veh) and reduced or rescued in in SST-Control+GL-II-73 and SST-Silenced+GL-II-73 mice (*p*<.01; **Fig 4*n***). Analysis of Z-memory impairment (YM, NORT) revealed a main effect of group (*F_1,41_*=15.18; *p*<.001) and trend-level group*treatment interaction (*F_1,41_*=3.29; *p*=.07), wherein SST+ cell silencing increased scores (*p*<.05 vs. SST-control+Veh) and GL-II-73 partially rescued these (*p*=.059; **Fig 4*o***). There was no effect of sex for all Z-scores.

## Discussion

In this report, we validated an approach to manipulate brain-wide SST+ cell function by selectively expressing hM4Di DREADD via i.c.v. infusion of AAV-PHP.eB in neonatal *Sst^Cre^* mice. In adulthood, brain-wide SST+ cell silencing increased neuronal activity in the PFC, HPC, and amygdala, elevated plasma corticosterone, and increased anxiety- and anhedonia-like behaviors, behavioral emotionality, and memory impairment. Treatment with GL-II-73, an α5-GABAA-R-selective PAM, rescued each of these behavioral changes. These results independently confirm prior evidence that SST+ cells regulate mood and cognitive functions, suggesting that their disruption may contribute to MDD-related symptom dimensions, and show for the first time that augmenting α5-GABAA-R function can rescue behavioral deficits induced directly by reduced SST+ cell function.

### Silencing brain-wide SST+ cell activity

To selectively silence brain-wide SST+ cell function, we employed neonatal i.c.v. infusion of FLEx-ed AAV-PhP.eB hM4Di vectors in *Sst*^Cre/+^ mice. AAV-PHP.eB capsids have enhanced GABAergic interneuron transduction (Gholizadeh et al., 2013), but previously relied on high-dose systemic administration (Deverman et al., 2016; Chan et al., 2017). Here, we validated high SST+ cell transduction via low-dose i.c.v. infusion in neonates (Kim et al., 2013) as a cost-effective technique overcoming transgenic colony maintenance and genetic leakage that is sometimes reported in lox-stop-lox models (Madisen et al., 2012). In *Sst^Gfp/+^* mice, cell specificity (~98-99%) and efficiency (56-95%) were high, but not complete, consistent with systemic AAV-PHP.eB in *Sst^Cre/+^* mice (Allen et al., 2017). This may be due to lower IHC detectability in *Sst^Cre^* mice that have reduced *Sst* levels and contain small numbers of non-SST Cre- or GFP-expressing cells (Ma et al., 2006; Taniguchi et al., 2011; Hu et al., 2013; Viollet et al., 2017). Of note, high but incomplete SST+ cell silencing may more closely reflect the low (but not ablated) cellular phenotype observed in human MDD.

Whereas past approaches used region-specific cell knockdown or *Sst* deletion (Soumier and Sibille, 2014; Lin and Sibille, 2015), our approach silenced intact SST+ cells, decreasing both GABA and SST release. These changes may be more closely related to circuit changes in MDD, where an intrinsic vulnerability causes low SST and GABA markers per cell and across corticolimbic brain regions (Fee et al., 2017). SST+ cell silencing was validated by increased neuronal activity in the PFC, HPC, and Amy in the absence of a stimulus, implying inhibitory neuron silencing and consistent with past approaches (Soumier and Sibille, 2014; Allen et al., 2017). SST+ cell targeting was validated via qualitative inspection throughout the neocortex, and confirmed in all mice. Our approach did not distinguish SST vs. GABA roles, modeling evidence of both markers being reduced in MDD (Fee et al., 2017). However, corticolimbic hyperactivation is an expected outcome reflecting silencing both mediators given their shared roles in PN inhibition (Tallent and Siggins, 1997; Schweitzer et al., 1998; Stengel and Taché, 2017), and consistent with past chemogenetic SST+ cell silencing studies (Soumier and Sibille, 2014; Allen et al., 2017) and chronic stress studies wherein GABA and SST markers are selectively reduced (Lin and Sibille, 2015; Girgenti et al., 2019; Fee et al., 2020). Plasma corticosterone elevation also validated reduced SST function, given its role in inhibitory regulation of corticosteroid release (Engin and Treit, 2009; Prévôt et al., 2016, 2018). That corticosterone effects only reached trend level may reflect the long measurement window chosen (i.e., designed for c-Fos analyses); indeed *Sst* ablated mice had elevated corticosterone at baseline, but not during or after stress (Lin and Sibille, 2015). Behavioral findings were also consistent with mice having low GABAA-R signaling (Ren et al., 2016) or SST knockdown (Soumier and Sibille, 2014; Lin and Sibille, 2015). Given that GABA-AR-acting BZDs and SST receptor-acting analogs confer anxiolytic- and antidepressant-like effects in rodents (Sanders and Shekhar, 1995; Engin et al., 2008; Yeung et al., 2011; Prévôt et al., 2016), evidence suggests that both systems regulate mood and cognitive functions.

Behavioral deficits were induced sub-chronically (i.e., repeated acute exposure across 9 tests), and were relatively lasting (i.e., detected 6 hours after CNO injection in PT), consistent with past chemogenetic studies (Guettier et al., 2009; Alexander et al., 2010). The CNO dosage was selected based on brain penetrant levels >EC_50_ for hM4Di from ~15-30+ minutes after administration and clearing from plasma within two hours (Jendryka et al., 2019). Thus, behavioral deficits likely reflect direct consequences of reduced SST+ cell regulation of corticolimbic functions.

The DREADD approach has several technical caveats, including CNO to clozapine back-metabolism that may confer off-target effects (Gomez et al., 2017; Jendryka et al., 2019). However, CNO-derived clozapine did not exceed hM4Di EC_50_ at the selected dose (Gomez et al., 2017), and CNO alone did not alter behavior in *Sst^Cre/+^* control mice (**Fig S1**) or locomotion in *Ss1^hSyn-hM4Di-mCherry^* mice (**Fig 3**). Furthermore, when all groups received CNO, behavioral changes were detected only in hM4Di-expressing mice (**Fig 4**). As human pathologies reflect chronic conditions, we tested chronic SST+ cell silencing via repeated i.p. injections or oral CNO delivery but found no behavioral changes (*unpublished data*). Rather than this being a consequence of chronic SST+ cell silencing, we attributed these effects to technical limitations of DREADDs, including that CNO has a rapid half-life and is challenging to administer at levels exceeding hM4Di EC_50_ orally over long time-periods, without back-metabolism and accumulation of lipophilic clozapine (Gomez et al., 2017; Jendryka et al., 2019). Indeed, evidence of chronic chemogenetic hM4Di activation is sparse, including null or opposite effects compared to acute hM4Di activation (Soumier and Sibille, 2014; Carvalho Poyraz et al., 2016; Urban et al., 2016). Conversely, chronic excitatory DREADD activation has been achieved using larger CNO doses and due to the unique pharmacokinetic properties of hM3Dq (Jain et al., 2013). These challenges justify the ongoing development of improved DREADD actuators or delivery systems (Bonaventura et al., 2019).

### SST+ cells and depressive-like behavior

Low SST and GABA markers in postmortem MDD subjects, and depressive-like deficits in mice with altered SST+ cell function, suggest a contributing role in MDD (Fee et al., 2017). We replicated past findings, including anxiety-like behaviors (OFT, EPM) observed in *Sst* knockout (Lin and Sibille, 2015) and acute chemogenetic PFC SST+ cell-silenced mice (Soumier and Sibille, 2014). Our findings extended roles for SST+ cell regulation of behavioral deficits to antidepressant-predictive (e.g., FST) and short-term memory dimensions (e.g., NORT). However, we did not find working memory impairment as with optogenetic PFC SST + cell silencing, possibly due to distinct inhibition timing, i.e., initiated before vs. during the test (Abbas et al., 2018) or anxiogenic reactivity of *Sst^Cre^* mice to frequent experimenter handling (Viollet et al., 2017). Notably, past studies found that acute and chronic blockade of PFC SST+ cell function had opposite physiological and anxiogenic outcomes (Seybold et al., 2012; Soumier and Sibille, 2014). However, region-specific manipulations may confer compensatory neural circuit adaptations that are not reflective of human pathology, wherein reductions in SST+ cell markers are evident across multiple brain regions in human and animal studies (Fee et al., 2017). Indeed, we found behavioral changes that closely paralleled past rodent chronic stress studies (Nikolova et al., 2018; Prevot et al., 2019c; Fee et al., 2020) that also found selective impingement upon SST+ cell functions via dysregulated cell integrity pathways (Lin and Sibille, 2015; Girgenti et al., 2019; Oh et al., 2019). These findings suggest that SST+ cell deficits are an intermediary causal factor between upstream risk factors (e.g., chronic stress, altered proteostasis and neurotrophic factor signaling) and MDD symptom emergence (Fee et al., 2017; Prevot and Sibille, 2020).

In SST-control vs. SST-silenced mice (using control vs. hM4Di viruses), we replicated anxiogenic findings from the first characterization experiment (using veh vs. CNO), but found stronger loading on anhedonia-like deficits. Given that anxiogenic changes were consistent in the PT, a test showing enhanced reproducibility and reliability compared to classical tests (Nikolova et al., 2018; Prevot et al., 2019c), inter-cohort differences may be due to the variability of classical tests that were designed to detect pharmacological treatment effects rather than disease-related processes (Willner, 2017; Prevot et al., 2019c).

Given that SST+ cell deficits are more severe in females (Seney et al., 2013; Seney, M., Tripp, A., McCune, S., Lewis, D., Sibille, 2015), we included sex as a covariate. However, differences were sparse as, for example, we identified increased anhedonia-like behavior in female mice overall that was unrelated to SST+ cell silencing.

### Antidepressant potential of α5-PAM

α5-GABAA-R potentiation via GL-II-73 had antidepressant-, anxiolytic-like, and pro-memory effects in mice with brain-wide low SST+ cell function. GL-II-73 rescued deficits in the EPM and FST, consistent with effects in wildtype mice (Prevot et al., 2019b) and with phenotypes of mice having disinhibited SST+ cell function (Fuchs et al., 2016). However, CNO+GL-II-73 induced hyperlocomotor effects in the EPM, OFT, and presumably the mobility-dependent FST (**Table S2**), so these results were excluded from Z-score analysis. These changes did not appear in past studies of GL-II-73 alone (Prevot et al., 2019b), so we cannot preclude potential drug-drug interactions. However, upon exclusion of these tests, Z-scoring revealed that GL-II-73 reduced anxiety-, anhedonia-like behavior, overall emotionality and (trend-level) memory impairment in both SST-control and SST-silenced mice, consistent with past studies (Prevot et al., 2019b). Contrasting findings in chronic stress-exposed mice, GL-II-73 did not improve YM performance, potentially due to increased *Sst^Cre^* mice reactivity to experimenter handling necessary for this test (Viollet et al., 2017). Indeed, SST-controls had lower YM alternation rates compared to wildtype mice in past studies (Prevot et al., 2019b). Further, GL-II-73 induced trend-level rescue of NORT deficits (a low-handling test), consistent with HPC α5-GABA-AR mediation of short-term memory (Möhler and Rudolph, 2017).

Others have reported antidepressant-like and pro-cognitive properties from α5-knockdown or negative allosteric modulators (NAMs) (Martin et al., 2010; Fischell et al., 2015; Zanos et al., 2017; Bugay et al., 2020). Although these findings appear to conflict with α5-PAMs, it may be that SST+ cell regulation of microcircuit activity benefits from “tightening” in contexts characterized by under-inhibition (i.e., poor information encoding) and “loosening” in contexts characterized by over-inhibition (i.e., poor information transfer), supporting the utility of α5-PAM/NAM for different contexts, behaviors, or disorders (Prevot and Sibille, 2020). Indeed, we found that α5-PAM had therapeutic effects in SST-silenced mice with corticolimbic hyperactivity. Another possibility is that α5-PAMs and NAMs could increase the signal-to-noise ratio of corticolimbic microcircuits through different mechanisms, for example by promoting phasic vs. tonic PN activity with α5-PAMs or by normalizing PN signaling through ketamine-like synaptic plasticity with α5-NAMs (Bugay et al., 2020).

### Future directions

Despite marked improvement in sedative and amnesic side-effect profiles of BZDs, α5-selective derivatives such as GL-II-73 have mild α1-3 GABAA-R potentiation (Prevot et al., 2019b) that may worsen or improve therapeutic efficacy. For example, more selective α5-PAMs (Stamenić et al., 2016) had weaker antidepressant-like and pro-cognitive effects only in chronic stress-exposed female (Piantadosi et al., 2016) and non-stressed male mice (Prevot et al., 2019b). Given that regional localization impacts α1-5-subunit functional differentiation (Sieghart and Sperk, 2002), characterization studies using gene profiling (Lin and Sibille, 2015) or RNA labeling (Hu et al., 2019) may inform therapeutic strategies.

In conclusion, we demonstrated that brain-wide SST+ cell function regulates mood and cognitive functions and found support that acute disruptions contributing to anxiety- and anhedonia-like behaviors, overall behavioral emotionality, and impaired memory may reflect similar processes in psychiatric diseases over a longer scale. Deficits arising from brain-wide low SST+ cell function were rescued by α5-PAM, representing a promising new avenue for the development of targeted antidepressants.

## Acknowledgements

We acknowledge the hard work of CAMH institutional animal facility staff for their assistance in breeding, genotyping, and maintaining colonies. Special thanks to K.F., G.F., and K.D. Special thanks to Dr. Bryan Roth for providing DREADD vectors (Addgene #44362) and Drs. Gradinaru and Deverman for the AAV-PHP.eB serotype technology. Thanks also to the Penn Vector Core for packaging and production of AAV-PHP.eB serotype control viruses (Addgene #50459).

## Author Contributions

C.F., M.B., and E.S. conceived of the study design. C.F. performed all experiments, data acquisition, and analysis with assistance from T.P., K.M., and M.B. C.F. wrote the manuscript with critical input from T.P., M.B., and E.S. D.K., G.L., P.M., and J.C. contributed to distinct steps for the synthesis and development of GL-II-73.

