## Supplementary Table and Figures for "Depressive-like behaviors induced by somatostatin-positive GABA neuron silencing are rescued by alpha 5 GABA-A receptor potentiation"

*Corresponding author

**Running title**: SST+ Cell silencing and depressive-like deficits

**Contents**

| **Supplementary Materials and Methods** | 3 |
| --- | --- |
| Model Development | 3 |
| Behavioral Assessment | 5 |
| Corticosterone and Immunohistochemistry Analysis | 8 |
| Data Analysis | 9 |
| **Supplementary Tables** | 10 |
| **Supplementary Table S1.** Characterization experiment 1 secondary test parameters for locomotor activity, home cage readouts, fluid intake, pre-trials, and significant sex effects | 10 |
| **Supplementary Table S2**. Characterization experiment 2 secondary test parameters for locomotor activity, home cage readouts, fluid intake, pre-trials, plus significant sex effects | 11 |
| **Supplementary Figures** |  |
| **Supplementary Figure S1.** Lack of anxiety-like behavior or locomotor alterations from 3.5mg/kg clozapine-N-oxide (CNO) in *Sst^Cre^* mice | 13 |

**Materials and Methods**

**Model Development**

**Animals.** To overcome low immunohistochemistry (IHC) signal in *Sst^Cre^* knock-in mice that have lowered SST expression [34], cell type specificity of viral transduction was validated in mice expressing green fluorescent protein (GFP) in SST+ Cells using the *Sst^Gfp^* mouse line, as described previously [33,48]. Briefly, mice were crossed with *Sst* promoter-directed Cre recombinase expression (*Sst^tm2.1(cre)Zjh^*/J, Jackson Laboratories, Bar Harbor, ME; #913944) with mice carrying a loxP-flanked STOP cassette controlling GFP expression (Ai6 Rosa26-loxP-STOP-loxP-ZsGreen; Jackson Laboratories; #008242).

Behavioral and molecular experiments were performed in heterozygous *Sst^Cre/+^* mice (*Sst^tm2.1(cre)Zjh^*/J, Jackson Laboratories, Bar Harbor, ME; #913944) on a C57BL/6J background receiving viral vectors (see below). All mice were postnatal day (PND) 0 at surgery and 9-16 weeks at testing. Mice were maintained under a 12-hour light/dark cycle with food and water *ad libitum*, group-housed (4/cage), except during behavioral testing (1/cage). All tests were performed in accordance with Institutional and Canadian Council on Animal Care (CCAC) guidelines.

**AAV Vectors.** Enhanced serotype inhibitory DREADD vectors (AAV-PHP.eB-hSyn-DIO-hM4D(Gi)-mCherry; Addgene, Cambridge, ME, #44362) were a gift from Dr. Bryan Roth using technology from Drs. Gradinaru and Deverman [49]. The Control vector (AAV-PHP.eB-hSyn-DIO-mCherry; Addgene #50459) was produced by Penn Vector Core (Philadelphia, PA).

**Neonatal Viral Vector Injection.** Brain-wide SST+ Cell-specific hM4Di expression was achieved by bilateral ventricular infusion of low-dose PHP.eB serotype Flip-Excision (FLEx-ed) AAVs in neonatal *Sst^Cre/+^* mice, as previously described [50]. In brief, neonates were collected from the dam within 12 hours of birth, cryoanesthetized on ice, and received bilateral AAV injections (1 µl/side, all experiments) using a 5 µl Hamilton syringe with 32G needle (Reno, NV; #7634-01). Injections targeted 3 mm depth perpendicular to a site equidistant between lambda and bregma, ±1 mm lateral to midline [50]. Neonates recovered on a heating pad and were returned to mothers with minimal disturbance until weaning. For validation experiments, *Sst^GFP^* neonates received ~10^13^ vg/mL AAV-PHP.eB-hSyn-DIO-hM4D(Gi)-mCherry virus. For characterization experiments, *Sst^Cre/+^* neonates received ~10^13^ vg/mL AAV-PHP.eB-hSyn-DIO-hM4D(Gi)-mCherry virus (*Sst^hSyn-hM4Di-mCherry^* mice) or AAV-PHP.eB-hSyn-DIO-mCherry virus (*Sst^hSyn-mCherry^* mice).

**Chemogenetic Inhibition of Brain-wide SST+ Cells & Co-administration with α5-PAM.** In the first experiment, adult *Sst^hSyn-hM4Di-mCherry^* mice received clozapine-N-oxide (CNO; 3.5 mg/kg) or vehicle (Veh; 0.9% saline) intraperitoneally (i.p.) 30 min before testing (*n=*16/group; 50% female). CNO dosage was based on a study reporting brain penetrant levels >EC_50_ for hM4Di in mutant mice, with no effect in non-hM4Di mice [51]. To preclude off-target effects, we assessed locomotor activity and anxiety-like behavior in separate cohorts of *Sst^Cre/+^* administered Veh or CNO, finding no difference between groups (**Figure S1**).

In the second experiment, we assessed whether the deficits induced by SST+ Cell silencing could be rescued by co-administering a validated α5-PAM, GL-II-73, with CNO [43]. *Sst^hSyn-mCherry^* and *Sst^hSyn-hM4Di-mCherry^* mice were generated by neonatal infusion of the control or inhibitory DREADD virus, respectively. All groups were administered CNO (3.5 mg/kg i.p,), thus achieving SST silencing only in hM4Di-expressing mice (SST-silenced vs. SST-control groups). To test the effects of GL-II-73, *Sst^hSyn-mCherry^* and *Sst^hSyn-hM4Di-mCherry^* mice were randomly assigned to receive GL-II-73 (10mg/kg diluted in 85% dH2O, 14% propylene-glycol, and 1% Tween-80, plus CNO i.p.), or vehicle only (Veh, plus CNO i.p.) (4 groups, *n*=10-12/group; 50% female) [52].

**Behavioral Assessment**

Behavioral testing commenced in adulthood (9 weeks), assessing *anxiety-like behavior* with the phenotyper test (PT), elevated plus-maze (EPM), open field test (OFT) and novelty-suppressed feeding test (NSF), *anhedonia-like behavior* with sucrose consumption test (SCT), and *mixed anxiety-/anhedonia-like* *behavior* with novelty-induced hypophagia test (NIH), *antidepressant-predictive behaviors* in the forced-swim test (FST), and *memory impairment* in the Y-maze (YM) and novel object recognition test (NORT), following past experimental design [51]. All tests were conducted under blinded conditions during the light cycle 30 min following drug treatment, except for the dark-cycle PT where drugs are administered 1 hour before light challenge. Sample sizes varied up to ±5 mice per test due to exclusions as a result of errors in tracking or recording (PT, EPM, OFT, NORT), food/water deprivation (NSF, NIH, SCT), or apparatus failures (YM, EPM); however exclusions were balanced across groups.

**Phenotyper Test**. Our lab developed the PT to measure anxiogenic light challenge response in a home cage-like environment [52]. Ethovision software tracks time spent in the PhenoTyper^TM^ apparatus (Noldus, Leesburg, VA) with designated food and shelter zones overnight throughout the dark cycle (7pm-7am). In the short protocol (adjusted for drug half-life), mice are habituated to the arena at 5pm, injected at 6:30pm prior to the dark cycle. 1 hour after dark cycle start, a spotlight illuminates the food zone (8-9pm), inducing a characteristic avoidance pattern in mice (i.e., seeking shelter until the light turns off) that is protracted up to 5 hours in mice exposed to chronic stress [52]. The extent of anxiogenic response was measured throughout testing and summarized by shelter zone area under the curve (AUC): the summed averages of 30 minute bins from the initiation of light challenge until the end of the test.

**Elevated Plus Maze.** The EPM was situated 55cm from the floor with 4 white Plexiglas arms, with two open (27x5cm) and two enclosed arms (27x5x15cm) located parallel. 30 minutes after Veh/CNO injection, animals explored the EPM for 10 min in a dimly lit room with an aerial-mounted camera. Time spent (s) in open arms and distance travelled (m) were measured Using ANY-maze software.

**Open Field Test.** 30 minutes after Veh/CNO injection, mice freely explored an open arena (70x70x33cm) for 10 min. Video tracking using ANY-maze software divided the field into 2 concentric squares of equal area (1633cm^2^), measuring time spent in the inner zone (s) and distance travelled (m).

**Novelty-suppressed Feeding**. In the NSF, mice were food deprived for 16 hours. Deprivation was controlled by bedding change 48 hours prior, food inspection and removal, and weighing before and after. Food-deprived mice were tested in order by percentage weight loss (greatest to least), and 30 min after weight/volume-adjusted injection of Veh/CNO, underwent a timed trial for latency to feed on a food pellet placed in the middle of a brightly-lit novel arena (62x31x48cm, 12 minutes maximum). As a control for appetitive drive, latency to feed was also measured in the home cage, immediately following the novel environment test (6 minutes maximum).

**Novelty-induced Hypophagia.** Mice were habituated to a liquid reward (1mL of sweetened condensed milk) for 2 days, then injected with Veh/CNO on day 3 and 30 minutes later tested for latency to drink the milk in the home cage (to control for appetitive drive). On day 4, 30 minutes after Veh/CNO injection, latency to drink milk was assessed in a bright-lit new cage (26x15x12cm, 6 minutes maximum).

**Sucrose Consumption Test**. Mice were habituated to 1% sucrose solution for 48 hours, fluid deprived for 14 hours, injected with Veh/CNO and 30 minutes later assessed for intake (mL) over 1-hour re-exposure. Mice were returned to water for ~2 days, and the same procedure was repeated, including Veh/CNO injection to assess water intake (“water consumption test”). Sucrose consumption volume was used as a primary measure, and sucrose ratio [sucrose/(sucrose + water consumed)] was used as a secondary measure to control for any differences in thirst.

**Forced Swim Test.** 30 minutes after Veh/CNO injection, mice were placed in a water-filled beaker (25 cm height, 24±1^o^C) and recorded during a 6 min swim trial. Immobility, defined as the minimum amount of movement to stay afloat, was manually tracked from videos by an experimenter blind to treatment.

**Y-Maze.** Mice underwent habituation to the YM apparatus (3 equally-spaced 26x8x13 cm arms) and distal cues across 10 min free exploration for 2 days. On day 3, mice were trained to alternate between arms across 7 trials, where they are closed in the arm of their choosing for 30s, separated by 30s inter-trial interval (ITI) in the start box. On day 4, mice were administered Veh/CNO and 30 minutes later tested on 7 successive trials with a 90s ITI. An alternation rate was calculated measuring successful arm alternations (of 6 possible). To preclude motivation bias, an 8^th^ trial with 5s ITI was added wherein mice failing this trial were excluded from analyses.

**Object Recognition.**  On consecutive days, mice were habituated to an open field (70x70x33 cm), first empty and then with 2 identical plastic objects serving as “familiar” objects (10 min/day). On day 3, mice explored the familiar objects for 10 min, then underwent a 3 hour delay before a second 10 min trial (preceded 30 minutes by Veh/CNO treatment) where they explored one familiar object and one “novel” object of similar size. A discrimination ratio of novel object preference [novel time/(novel + familiar time)] measured short-term memory for the familiar object, given rodent preference for novelty [53]. A pre-trial discrimination ratio [familiar_1_/ (familiar_1_ + familiar_2_)] was calculated to preclude arena side preference.

**Behavioral Z-Score Summaries**

To assess consistency across related behavioral tests, we used z-score normalization to capture changes along behavioral dimensions, including anxiety-like (PT, EPM, OFT, NSF, NIH), anhedonia-like (NIH, SCT), and overall behavioral emotionality (all previous + FST), or memory impairment including working- and short-term memory (YM, NORT) [54]. This approach summarizes inter-related behavioral parameters, as validated in multiple preclinical models [35,42,52,55]. Test parameters were normalized to the mean and standard deviation of control groups and expressed in a uni-directional manner relative to controls, i.e., increased Z-score = deficit and decreased Z-score = improvement. Normalized z-scores for each test were averaged across tests within each dimension to capture overall deficit scores (e.g., Z-anxiety).

**Corticosterone Measurement**

To determine the effect of SST+ Cell silencing on basal corticosterone levels, *Sst^hSyn-hM4Di-mCherry^* mice from behavioral experiments were injected with Veh or CNO 110 min prior to blood collection via cardiac puncture under deep anesthesia (250 mg/kg Avertin i.p.) before perfusion. Blood samples were collected between 10:00-12:00 in heparinized tubes and centrifuged at 4^o^C, 10,000 RPM for 10 min to isolate plasma. Plasma corticosterone was assayed using a commercial enzyme-linked immunosorbent assay (ELISA) kit (Arbor Assays, Ann Arbor, MI). All samples were run in duplicate according to the manufacturer’s protocol.

**Immunohistochemistry and Microscopy**

We first assessed DREADD virus efficiency and specificity in *Sst^Gfp^* mice. *Sst^Gfp^* mice were perfused with 4% paraformaldehyde, brains were then harvested and cryosectioned at 40 μm using a cryostat (Leica Microsystems, Concord, ON). Sections were double-stained with goat anti-GFP (1:1000, Rockland, Limerick, PA; #600-101-215) and chicken anti-mCherry (1:500, Abcam, Cambridge, MA; #205402), followed by donkey anti-goat Alexa Fluor^(R)^ 488 conjugate (1:500, ThermoFisher, Waltham, MA; #A-11055) and donkey anti-chicken Rhodamine Red^TM^-X conjugate (1:500, Jackson Immunoresearch, West Grove, PA; #703-295-155). Images were acquired using an Olympus IX83 inverted microscope (Richmond Hill, ON, Canada) with Hamamatsu ORCA-Flash4.0 camera (Bridgewater, NJ, USA) and ProScan-III motorized stage (Prior Scientific, Rockland, MA, USA). DREADD virus transduction efficiency and SST+ cell specificity was assessed via quantification of overlap between mCherry and GFP, for efficiency (mCherry/GFP) and specificity (GFP/mCherry) (*n*=8 mice/group, 2 fields of view/region/mouse and summed as an average across all fields for “overall”).

CNO-hM4Di binding increases activation of inward rectifying potassium channels and decreases cAMP signaling, ultimately causing transient membrane hyperpolarization [55,56]. Thus, c-Fos+ cell counts served as a marker to validate local neuronal activation following reduced inhibition due to SST+ Cell silencing [33,56,57]. Two days after the last behavior test, *Sst^hSyn-hM4Di-mCherry^* mice were treated with vehicle/CNO 110 min prior to blood collection/perfusion [51,56]. Cryo-sections were immunostained with rabbit anti-c-Fos (1:200, Synaptic Systems, Germany; #226003), followed by donkey anti-rabbit Alexa Fluor^(R)^ 488 conjugate (1:500, Jackson Immunoresearch; #703-295-155). c-Fos+ cells were counted in every third section totaling 4-5 sections/region/mouse spanning the medial prefrontal cortex (mPFC), dorsal hippocampus (dHPC), or basolateral amygdala (BLA). Brain regions were delineated at 2X using a mouse brain atlas [58], and 3D image stacks (333x333x0.75 μm) were collected at 20X by systematic random sampling, totaling 15-20 fields/section for PFC and dHPC and 4-5 for BLA. Image processing and cell counting was performed manually for Veh or CNO mice (*n* = 8/group) using ImageJ Fiji software [59] under blinded conditions.

**Data Analysis**

Data were analyzed using SPSS (IBM, Armonk, NY) and expressed as mean ± standard error of the mean (SEM). For the first experiment, behavioral parameters and IHC/ELISA data were analyzed using one-way analysis of covariance (ANCOVA) with treatment (CNO vs. Veh) as independent variable and sex as covariate. In the second experiment, behavioral parameters were analyzed using two-way ANCOVA with group (SST-control vs. SST-silenced) and treatment (Veh vs. GL-II-73) as independent variables and sex as covariate. Significant effects were followed up using Bonferroni adjusted *post hoc* tests. Time-course parameters from the PT and FST were assessed by repeated-measures ANCOVA with Greenhouse-Geisser correction for parameters failing sphericity assumptions. Object recall in the NORT was calculated using a paired-sample *t* test comparing time spent with left/right identical objects or familiar/novel objects.

**Supplementary Tables**

**Supplementary Table S1.** Characterization experiment 1 secondary test parameters for locomotor activity, home cage readouts, fluid intake, pre-trials, plus significant sex effects

| **Experiment 1** | | | | | | | | | | | | | | | | | | | |
| --- | --- | --- | --- | --- | --- | --- | --- | --- | --- | --- | --- | --- | --- | --- | --- | --- | --- | --- | --- |
| ***Test*** *Parameter (unit)* | ***Significant effect(s)*** ****p< .05;**p<.01; ***p*<.001; # *p<.1* | ***SST-DREADD+Veh*** | | | | | | ***SST-DREADD+CNO*** | | | | | | **Totals** | | | | | |
|  |  | Male | | | Female | | | Male | | | Female | | | Male | | | Female | | |
|  |  | Mean ± SEM | | | Mean ± SEM | | | Mean ± SEM | | | Mean ± SEM | | | Mean ± SEM | | | Mean ± SEM | | |
| PT Dist Travelled AUC | Not significant | 21144.28 ± 2673.06 | | | | | | 19590.63 ± 1913.36 | | | | | | 20367.45 ± 1622.92 | | | | | |
| EPM Dist Travelled | Not significant | 24.71 ± .84 | | | | | | 23.34 ± 1.63 | | | | | | 24.07 ± .87 | | | | | |
| OFT Dist Travelled | Not significant | 55.59 ± 3.28 | | | | | | 57.16 ± 3.33 | | | | | | 56.37 ± 2.30 | | | | | |
| NSF Home Latency | Not significant | 28.07 ± 2.44 | | | | | | 32.63 ± 3.89 | | | | | | 30.5 ± 2.37 | | | | | |
| NIH Home Latency | Not significant | 24.71 ± 3.32 | | | | | | 20.88 ± 3.06 | | | | | | 22.67 ± 2.24 | | | | | |
| WCT Consumed | Main effect of treatment** | 1.54 ± 0.13 | | | | | | 1.14 ± 0.11 | | | | | | 1.34 ± 0.09 | | | | | |
| PCT_SCT consumed | Not significant | 75.49 ± 11.12 | | | | | | 87.89 ± 11.81 | | | | | | 81.89 ± 8.07 | | | | | |
| SCT Consumed (males/females) | Main effect of sex** | 1.6 | ± | 0.073 | 0.85 | ± | 0.25 | 1.11 | ± | 0.22 | 0.81 | ± | 0.19 | 1.36 | ± | 0.13 | 0.83 | ± | 0.15 |
| YM Pre-trial | Not significant | 4.31 ± 0.29 | | | | | | 3.92 ± 0.31 | | | | | | 4.12 ± 0.21 | | | | | |

**Supplementary Table S2**. Characterization experiment 2 secondary test parameters for locomotor activity, home cage readouts, fluid intake, pre-trials, plus significant sex effects

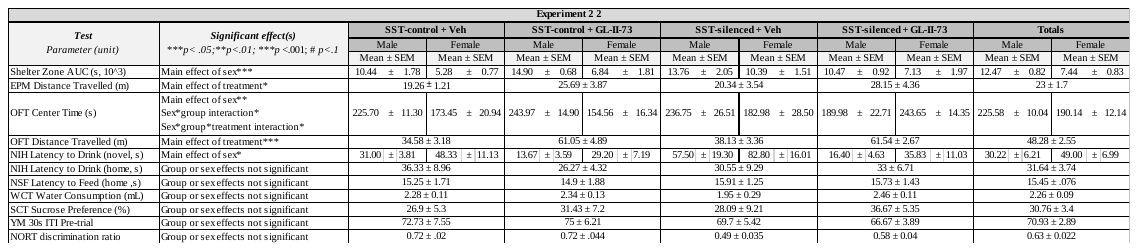

| PT Dist Travelled AUC (10^3^) | Not significant | 18197.28 ± 2397.94 | 23845.67 ± 2545.29 | 21393.2 ± 2648.59 | 21323 ± 3546.07 | 21253.46 ± 1403.24 |
| --- | --- | --- | --- | --- | --- | --- |

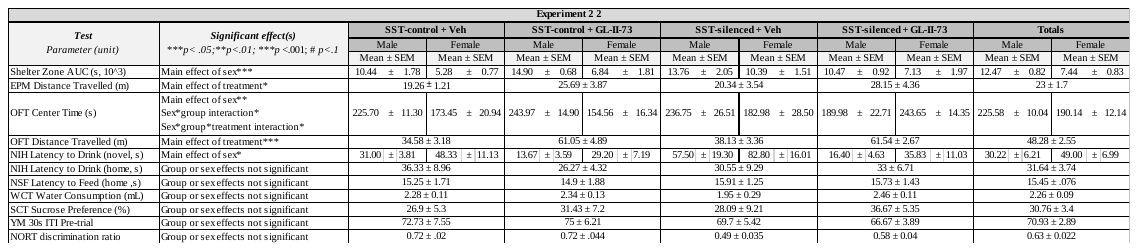

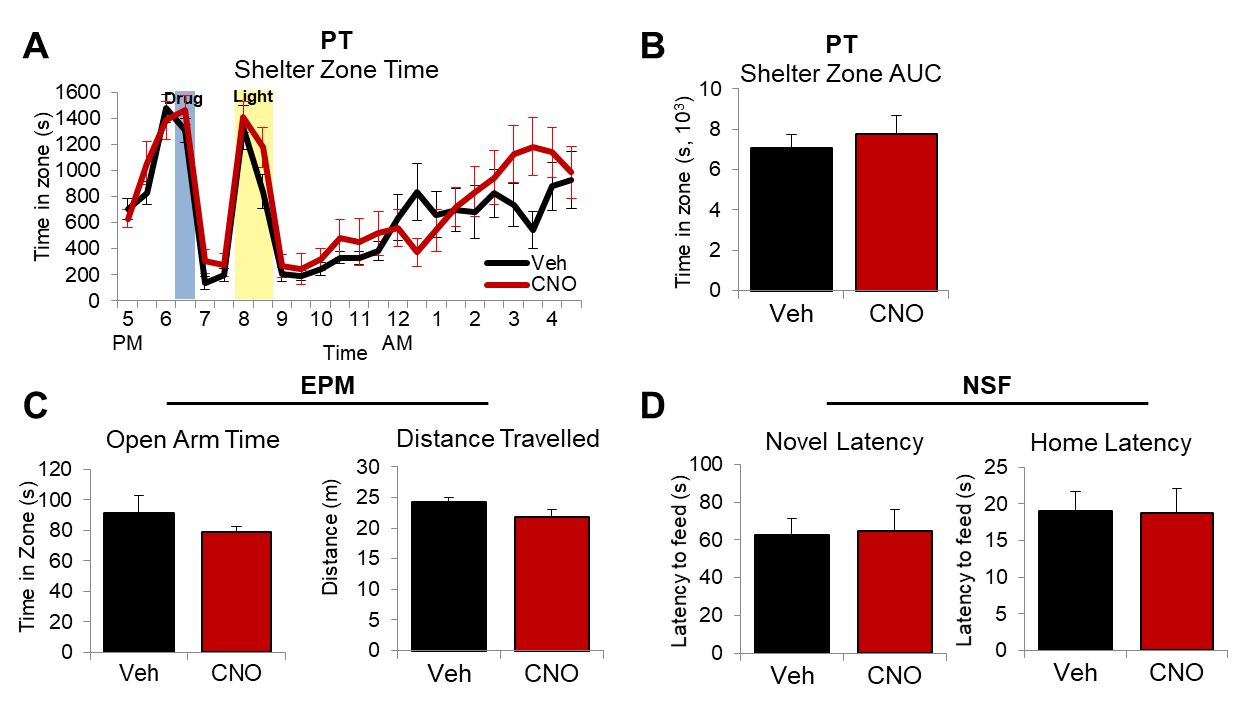

**Supplementary Figure S1.** **Lack of anxiety-like behavior or locomotor alterations from 3.5mg/kg clozapine-N-oxide (CNO) in *Sst^Cre^* mice** (*n =* 10-11; ~50% female) (B) Time course response to the PhenoTyper Test (PT) light challenge was not altered by CNO treatment (time: *F_7.31,138.98_* = 4.25; *p* < .001; treatment: *F_1,19_* = .98; *p* = .33; no interaction or sex effect). (B) Derived total shelter time area under the curve from PhenoTyper post-light challenge was not altered by CNO treatment (treatment: *F_1,20_* = 1.43; *p* = .25). (C) Time spent in the open arms (*F_1,18_* = 1.03; *p* =.32) and distance travelled (*F_1,18_* = 1.7; *p* =.21) in the elevated plus maze (EPM) was not significantly different between Vehicle (Veh)- and CNO-treated mice. (D) Latency to feed in a novel (*F_1,17_* = .19; *p* =.89) or home environment (*F_1,17_* = .08; *p* =.78) was not affected by CNO treatment in the novelty-suppressed feeding test (NSF).
